# A life-history axis of resource acquisition in bacteria: depolymerisation versus uptake

**DOI:** 10.64898/2026.10.25.761621

**Authors:** Yingyi Fu, Luciana Chavez Rodriguez, Josep Ramoneda, Katherine Shek, Jennifer B H Martiny, Steven D Allison, Yu Luo, Ashish A Malik

## Abstract

Microbial decomposition of complex organic matter is a major control on the global carbon cycle, and its efficiency hinges on how microbes acquire resources. Decomposition requires costly extracellular enzymes to break down complex polymers, whereas the resulting soluble products are captured far more cheaply through membrane transporters. How microbes balance investment between these two processes is a fundamental axis of microbial life history, yet it has remained difficult to quantify across their vast diversity. Using genomic repertoires for carbohydrate depolymerisation and uptake, we developed a simple ratio distinguishing depolymerisation-optimised from uptake-optimised strategies, and applied it to more than 249,000 environmental and host-associated bacterial genomes. These strategies were strongly phylogenetically structured and differed by habitat, with uptake-optimised bacteria unexpectedly more prevalent in diffusion-limited soils than in aquatic environments. In an 18-month field-scale plant litter decomposition experiment, uptake-optimised bacteria dominated during early decomposition, when fungi were more abundant and strongly expressed hydrolytic enzymes, shifting towards depolymerisation-optimised bacteria at later stages. A simple genomic ratio therefore captures ecologically meaningful resource-acquisition strategies that vary with phylogeny, habitat and successional stage. The balance of these strategies could shape community-level extracellular enzyme investment, with consequences for decomposition and carbon cycling.

## Main Text

Microorganisms, namely bacteria and fungi, drive the decomposition of complex organic matter and regulate the turnover and cycling of carbon^1,2^. Most polymers such as cellulose, hemicellulose, and lignin cannot be directly assimilated by microorganisms; they rely on extracellular depolymerisation to release soluble compounds that can subsequently be transported into cells^3,4^. At the community level, microbial resource acquisition traits linked to depolymerisation have been shown to reduce growth yield through a population-level trade-off: investment in extracellular enzyme production comes at the cost of cellular growth^5–7^. This reduction in growth yield, and therefore biomass, can affect carbon retention in soil, as much of soil organic matter is now recognised to be microbial in origin^8,9^. Microbial processing similarly governs carbon persistence in the ocean, where the microbial carbon pump converts labile organic matter into refractory compounds that persist over millennial timescales^10^.

In addition to extracellular depolymerisation of complex substrates, membrane transporter-mediated uptake of simpler compounds represents another fundamental component of microbial resource acquisition^6^. Although extracellular depolymerisation and membrane uptake can be intrinsically coupled during resource acquisition, microorganisms do not necessarily invest equally in these two processes^6,11^. Investment in extracellular enzymes increases access to complex substrates but incurs substantial metabolic costs in maintaining and expressing multi-gene depolymerisation systems and in secreting the enzymes, whereas investment in membrane transport enhances the capacity to capture available soluble compounds^6,12,13^. Variation in the relative investment in these functions may therefore give rise to differing resource acquisition strategies: a depolymerisation-optimised strategy that specialises in the extracellular breakdown of complex substrates, and an uptake-optimised strategy that focuses on the capture of soluble products. Because the uptake-optimised strategy incurs lower costs than the depolymerisation-optimised strategy, it is likely to entail smaller trade-offs with growth yield and biomass production.

Evidence that extracellular resource release and uptake can be decoupled has emerged from several microbial systems, particularly aquatic environments. An analogous asymmetry occurs in siderophore-mediated iron acquisition, where bacteria lacking siderophore biosynthetic pathways may retain transport systems for siderophore-bound iron^14^. Some marine pelagic bacteria consume polysaccharide hydrolysis products generated by neighbouring taxa despite lacking the extracellular hydrolytic enzymes required to release them^15^. Similar patterns have been observed during chitin degradation, where bacteria retain the machinery required for oligosaccharide uptake and metabolism while lacking the extracellular hydrolases needed to initiate polymer degradation^16^. Such cheater or exploiter behaviour may be widespread in natural communities, but it is difficult to demonstrate clear fitness cost to the depolymerising taxa. Collectively, these observations demonstrate that extracellular resource release and uptake capacities can be unequally represented across microorganisms, providing a mechanistic basis for contrasting depolymerisation- and uptake-optimised strategies.

Whether comparable resource acquisition strategies occur in soils, however, remains unclear. In contrast to relatively well-mixed aquatic environments, soils are characterised by tortuous pore networks, pronounced spatial heterogeneity and diffusion-limited movement of extracellular enzymes and their hydrolysis products, potentially allowing enzyme-producing microorganisms to retain a greater proportion of the benefits generated by their metabolic investment^17,18^. On this basis, one might expect soils to favour a depolymerisation-optimised strategy. Yet, soils also contain localised, transient pools of readily assimilable compounds generated by root exudation, microbial turnover, and organic matter decomposition^2,19^. These conditions could favour an uptake-optimised strategy which allows efficient capture of simple readily assimilable substrates within microscale hot spots. Interactions with neighbouring decomposers may further alter the relative costs and benefits of investing in depolymerisation versus uptake^17,20^. How these contrasting strategies are distributed across soil types and in comparison with other non-soil habitats remains poorly understood. Yet their balance could shape community-level investment in extracellular enzymes and, in turn, lead to variable rates of decomposition and the retention of carbon in different soils.

Beyond environmental variation, the relative investment in depolymerisation and uptake may also reflect evolutionary history and genomic constraints. Comparative genomic studies have shown that the potential to degrade structural polysaccharides is unevenly distributed across bacterial lineages, with closely related taxa often sharing similar repertoires of carbohydrate-degrading enzymes^21^. Transport capabilities likewise vary markedly among bacterial groups, including lineage-specific expansions of transporter families^22^. Together, these lineage-associated differences in depolymerisation and uptake machinery suggest that resource acquisition strategies may be phylogenetically structured^23^. These strategies may also be associated with broad genomic features. Maintaining diverse extracellular enzyme systems requires greater coding capacity, typically requiring larger genomes, and potentially higher metabolic costs, whereas genome streamlining can reduce investment in depolymerisation through gene loss while retaining transporters for the uptake of soluble compounds^24,25^. However, the link between these strategies and genome size remains unresolved.

The overall aim of this study was to develop and validate a genome-based framework to quantify bacterial resource acquisition strategies based on the balance between carbohydrate depolymerisation and uptake capacity. We examined if bacteria differ along a resource acquisition continuum with uptake-optimised bacteria having genomes with a greater repertoire of transporters for substrate uptake relative to extracellular depolymerisation enzymes, and depolymerisation-optimised bacteria having genomes with the opposite configuration. To test this framework, we analysed more than 249,000 environmental and host-associated metagenome-assembled genomes (MAGs) and characterised the global distribution of the two distinct resource acquisition strategies across habitats to test if uptake-optimised strategy is favoured in systems with higher diffusion rates and readily available substrates. We then focused on a collection of global soil MAGs to examine how these strategies vary across terrestrial biomes. We hypothesised that these contrasting strategies are phylogenetically conserved across the bacterial tree of life and that uptake-optimised bacteria would possess more streamlined genomes than depolymerisation-optimised bacteria owing to the loss or absence of multi-gene depolymerisation systems. Finally, we validated our genome-based framework using a field-scale plant litter decomposition experiment that tracked decomposition and succession of bacteria (and fungi) over 18 months. Together, these analyses provide an ecological basis for contrasting resource acquisition strategies in bacteria that could have differential implications for ecosystem processes such as carbon and nutrient cycling.

## Results and Discussion

### Contrasting resource-acquisition strategies across habitats

We first developed a simple ratio using genomic repertoires for carbohydrate depolymerisation and uptake to define an axis of bacterial resource acquisition and applied this genome-based framework to MAGs from environmental and host-associated microbiomes (n=249,311). Using microTrait^26^, we identified genes associated with complex carbohydrate depolymerisation and uptake from bacterial genome sequences (Data S1). Genes for complex carbohydrate depolymerisation comprised diverse glycoside hydrolase (GH) families acting on the major structural polysaccharides, including cellulose, xylan and other hemicelluloses, xyloglucan, mixed-linkage and other β-glucans, mannans and chitin. Genes for carbohydrate uptake comprised phosphotransferase systems (PTS), ATP-binding cassette (ABC) importers, major facilitator superfamily (MFS) permeases and outer-membrane sugar porins, mediating uptake of a broad range of mono-, di- and oligosaccharides and amino sugars. For each genome, these genes were converted into presence-absence profiles and summed within the two functional categories to estimate the breadth of depolymerisation and uptake potential. We then calculated the uptake-to-depolymerisation ratio as an index quantifying the contrasting strategies, with higher values indicating an uptake-optimised strategy and lower values a depolymerisation-optimised strategy (Fig. 1a).

**Fig. 1.**
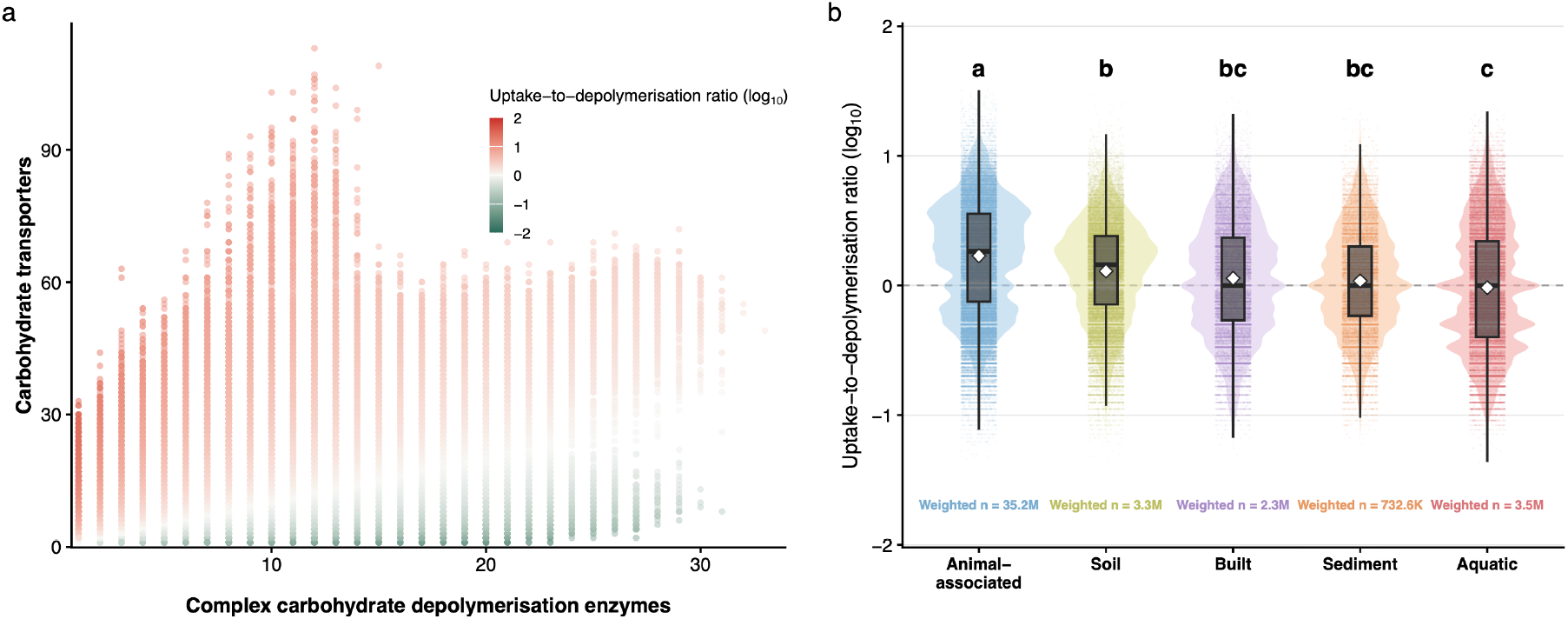
Contrasting microbial resource-acquisition strategies across environmental and host-associated microbiomes. (a) Relationship between the numbers of genes encoding carbohydrate uptake systems and complex carbohydrate depolymerisation enzymes across all genomes. Points are coloured by the uptake-to-depolymerisation ratio, illustrating the trait space defined by the two axes of bacterial resource acquisition. The ratio is log_10_-transformed, so a value of zero corresponds to a ratio of 1. Higher values indicate an uptake-optimised strategy (top left), whereas lower values indicate a depolymerisation-optimised strategy (bottom right). (b) Distribution of the uptake-to-depolymerisation ratio across major habitat types. Thick horizontal lines show the median, diamonds the mean, boxes the interquartile ranges, and whiskers extend to 1.5 times the interquartile range; violin plots show density distributions, with individual genomes overlaid as points. Each MAG was assigned to its dominant habitat based on its highest detection frequency; detection frequencies within that habitat were used as statistical weights. The weighted n represents the summed detection counts and does not correspond to the number of unique MAGs. Different letters indicate pairwise differences that were both statistically significant (least-squares model using heteroscedasticity-robust standard errors with Tukey adjusted *P* < 0.05) and of non-negligible magnitude (|Cliff’s δ| ≥ 0.147, corresponding to at least a small effect).

We then tested our first prediction that an uptake-optimised strategy would be favoured in higher-diffusion systems such as aquatic environments. Contrary to this expectation, uptake optimisation was instead lowest in these systems. MAGs from animal-associated microbiomes had the highest uptake-to-depolymerisation ratios followed by soil microbiomes with lower ratios in built, sediment and aquatic microbiomes in that order (Fig. 1b). The significantly higher ratios in soil than aquatic microbiomes, indicating a greater prevalence of uptake-optimised strategies, were unexpected, as this strategy is often assumed to be favoured in environments where soluble substrates diffuse more readily to cells. Although bulk diffusion is restricted in soils, soluble substrates are often generated locally within microscale hotspots through root exudation, microbial turnover and metabolic by-products^19,27,28^. This close spatial coupling between substrate release and cellular uptake may reduce the importance of long-distance diffusion and favour bacteria with greater uptake capacity^2^. Moreover, extracellular depolymerisation is metabolically costly, and some hydrolysis products may become accessible to neighbouring cells^29^. Thus, the high ratios observed in soil bacteria may reflect efficient capture of soluble compounds produced by nearby decomposers (a cheating behaviour) or supplied by root inputs and microbial turnover in proximity.

### Biogeographic patterns of soil bacterial resource acquisition strategies

Given the high prevalence of uptake-optimised bacterial genomes in microbiomes from soil, we next examined how the uptake-to-depolymerisation ratio varied across major terrestrial biomes using a separate database of global soil MAGs (n=28,932). We used the Whittaker biome classification based on mean annual precipitation and mean annual temperature to group MAGs into biomes of origin. MAGs from woodland/shrubland had the highest ratios, followed by tropical seasonal forest/savanna, tundra, temperate grassland/desert and subtropical desert (Fig. 2). Forests (temperate seasonal and boreal) had the lowest ratios. This pattern suggests that soil bacterial resource acquisition strategies vary along broad climatic gradients. The relatively greater uptake investment and lower depolymerisation investment in warmer, seasonally water-limited and less productive biomes than in colder, wetter systems could suggest an adaptation to increase efficiency of acquiring transiently available resources through uptake optimisation. On the contrary, higher prevalence of depolymerisation strategy in forests could be linked to higher availability of complex plant polymers from leaves and woody biomass.

**Fig. 2.**
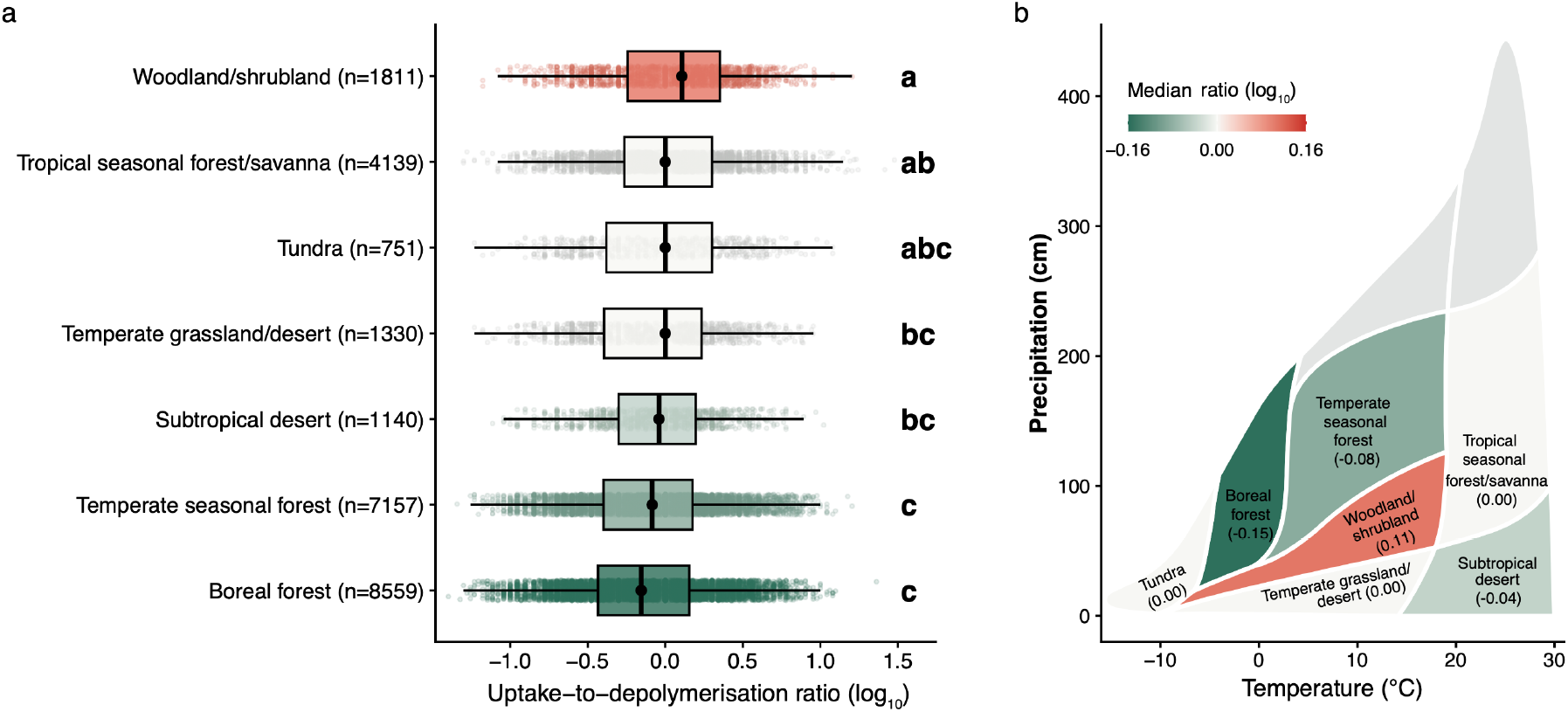
Biome-wide variation in bacterial resource-acquisition strategies across global soils. (a) Distribution of the log_10_-transformed uptake-to-depolymerisation ratio across major terrestrial biomes. Boxes show the median and interquartile range, whiskers extend to 1.5 times the interquartile range, and individual bacterial genomes are overlaid as points. Biomes are ordered by their medians. Different letters indicate pairwise differences that were both statistically significant (Dunn’s post hoc test with Benjamini–Hochberg correction following a significant Kruskal–Wallis test, adjusted *P* < 0.05) and of non-negligible magnitude (|Cliff’s δ| ≥ 0.147, corresponding to at least a small effect). Sample sizes indicate the number of genomes assigned to each biome. (b) Median uptake-to-depolymerisation ratio mapped onto Whittaker biome space.

The higher observed prevalence of uptake-optimised strategy in soils suggests that uptake optimisation may track proximity to readily available simple compounds, for example in the rhizosphere relative to bulk soil. We tested this prediction explicitly using subsets of the global soil MAGs comprising paired Poplar rhizosphere and bulk soils from three tree sites^30, 31^. MAGs from rhizosphere soil showed significantly higher uptake-to-depolymerisation ratio than those from adjacent bulk soil in two sites while the difference in the third site was marginal (Fig. S1). This indicates greater relative investment in carbohydrate uptake systems by rhizosphere bacteria relative to bulk soil bacteria to capture soluble substrates supplied through root exudation or turnover.

### Resource acquisition strategies exhibit clear phylogenetic structuring

To examine the evolutionary distribution of the two resource acquisition strategies, we mapped the uptake-to-depolymerisation ratio onto a reference bacterial phylogeny of MAGs from environmental and host-associated microbiomes (n=249,311). Uptake-optimised and depolymerisation-optimised genomes showed contrasting phylogenetic distributions across the bacterial tree of life. Actinomycetota, Cyanobacteriota, Bacillota, Pseudomonadota, and Chloroflexota generally exhibited higher uptake-to-depolymerisation ratios, whereas Bacteroidota, Patescibacteriota, Acidobacteriota, and Verrucomicrobiota had lower ratios (Fig. 3 and Fig. S2). In dryland-associated groups such as Actinomycetota, expanded transporter repertoires may favour rapid uptake of readily assimilable organic compounds released during transient pulses of resource availability, such as those following rewetting^32,33^. Several of these compounds are compatible solutes serving as both assimilable carbon sources and osmoprotectants under water stress. In Cyanobacteriota and some Chloroflexota lineages, high ratios may reflect autotrophic lifestyles (photoautotrophic or chemolithoautotrophic), which reduce reliance on extracellular depolymerisation for carbon acquisition^34,35^. The high ratios in Bacillota and Pseudomonadota are consistent with their reliance on versatile, high-affinity transport systems that favour rapid uptake of readily assimilable carbohydrates over investment in complex polysaccharides depolymerisation^36,37^. However, both phyla are metabolically diverse, and phylum-level averages mask substantial within-group variation, so the elevated mean ratio reflects a central tendency rather than a uniform strategy. The lower ratios in Bacteroidota, Acidobacteriota and Verrucomicrobiota are consistent with their well-documented investment in extracellular enzymes for depolymerisation of complex polysaccharides^38–40^. Bacteroidota in particular are characterised by polysaccharide utilisation loci (PULs) containing diverse CAZyme, or carbohydrate-active enzyme, repertoires for the breakdown of complex glycans^41^. The low ratio in Patescibacteriota likely reflects a different mechanistic basis: they are generally host-associated bacteria with streamlined genomes and encode few transporters rather than an expanded depolymerisation repertoire (Fig. S2), such that the low ratio may stem from low uptake potential rather than genuine decomposition capacity^42,43^.

**Fig. 3.**
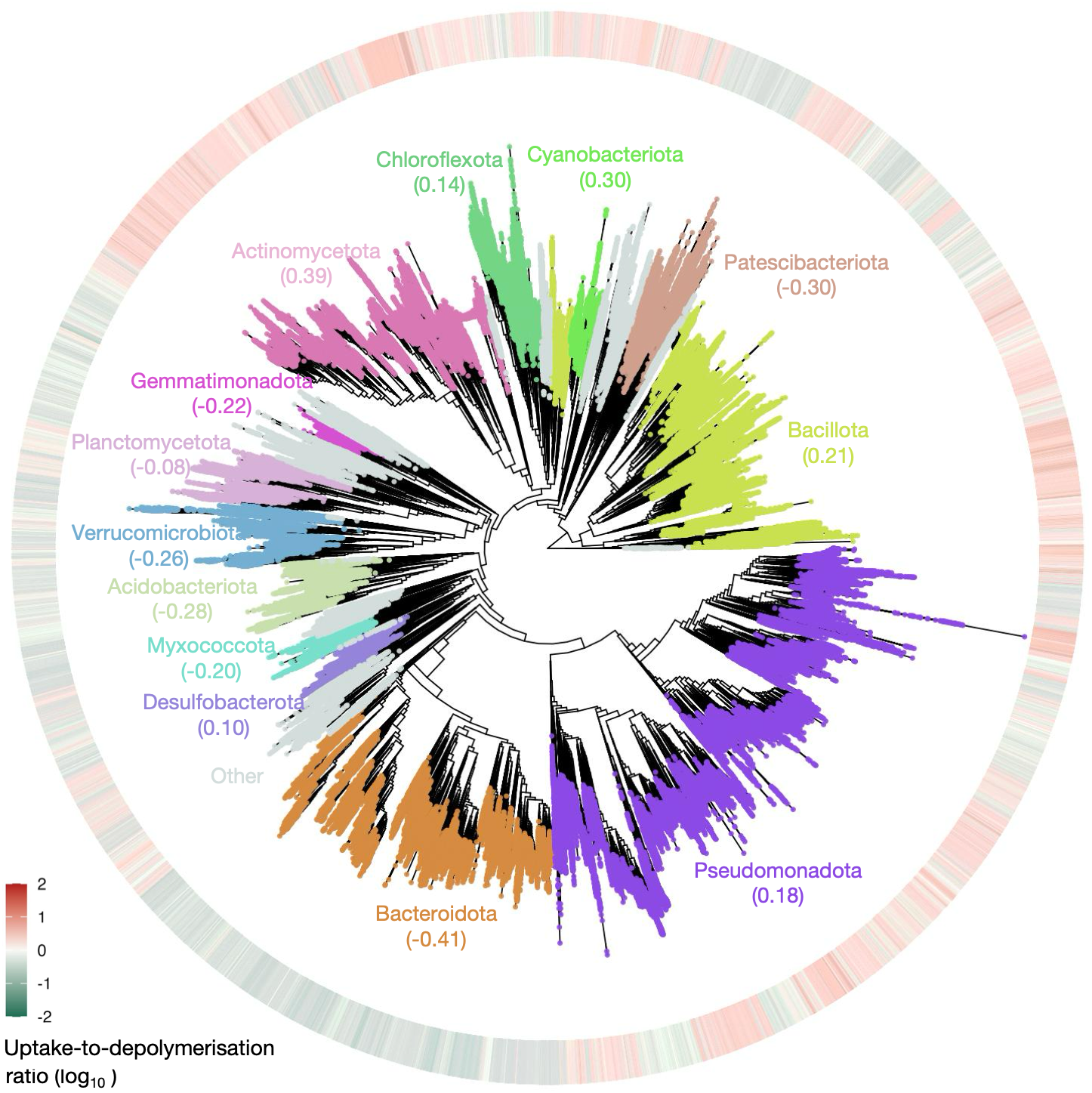
Phylogenetic distribution of bacterial resource acquisition strategies across global MAGs. A circular phylogenetic tree constructed from more than 200,000 environmental and host-associated bacterial MAGs. Major phyla are annotated around the tree, with their median log10-transformed uptake-to-depolymerisation ratio shown in parentheses. The outer continuous ring represents the log10-transformed uptake-todepolymerisation ratio for each MAG.

We next quantified the phylogenetic signal in bacterial resource-acquisition strategy by treating the log_10_-transformed uptake-to-depolymerisation ratio as a continuous trait and estimating Pagel’s λ across the bacterial phylogeny. The ratio exhibited a strong phylogenetic signal (λ = 0.956; *n* = 249,310; likelihood-ratio test against λ = 0, *P* < 0.001), indicating that closely related bacteria tended to have similar relative investments in uptake and depolymerisation. We also quantified the phylogenetic conservatism of uptake- and depolymerisation-optimisation strategies using consenTRAIT. Because consenTRAIT requires discrete rather than continuous traits, genomes in the top and bottom 10% of the uptake-to-depolymerisation ratio distribution were classified as uptake-optimised and depolymerisation-optimised, respectively. We found that depolymerisation-optimised bacteria exhibited significant phylogenetic conservation (τD=0.043, P<0.001), whereas uptake-optimised bacteria were more phylogenetically dispersed and did not differ significantly from a random expectation (τD=0.025, P=0.74). These results indicate that investment in extracellular depolymerisation is more strongly constrained by evolutionary history than in resource uptake. This pattern is consistent with previous observations that functionally complex traits, which depend on multiple coordinated genes, are generally more phylogenetically conserved than simpler traits such as resource uptake^23^. This may in part reflect that individual transporter genes are generally more readily gained by horizontal gene transfer than entire multi-gene depolymerisation systems^44^. More broadly, microbial functional traits commonly exhibit phylogenetic conservatism, although the strength of this signal varies among traits^23^. Our results extend these observations by showing that phylogenetic structuring applies not only to individual resource-acquisition traits but also to the relative allocation between substrate uptake and extracellular depolymerisation. Therefore, the position of bacteria along the uptake–depolymerisation axis represents a lineage-associated ecological strategy, potentially linking evolutionary history to differences in how bacterial communities access and process organic matter.

### Bacterial resource-acquisition strategies are largely decoupled from genome size

We then tested our hypothesis that uptake-optimised bacteria have streamlined genomes as an evolutionary consequence of losing or lacking multi-gene depolymerisation systems^45^. Across all genomes in the global dataset, the uptake-to-depolymerisation ratio showed only a weak, non-linear association with genome size (Fig. 4a). Genome size alone was therefore a poor predictor of bacterial resource-acquisition strategy, suggesting that the uptake-optimised strategy is largely decoupled from genome streamlining. However, the relationship between genome size and resource-acquisition strategy varied substantially among bacterial phyla (Fig. 4b-f), suggesting lineage-specific associations. The highest deviance was explained in Actinomycetota (Figure 4e), where the top 10% MAGs with highest ratios (uptake-optimised bacteria) had significantly smaller genomes than bottom 10% MAGs (depolymerisation-optimised bacteria), providing some evidence of genome streamlining in uptake-optimised bacteria of this phylum. Actinomycetota also had the highest median ratio (Fig. S2), and thus the highest prevalence of uptake-optimised strategy. Together, these results suggest that uptake optimisation may contribute to genome streamlining in some linages but is likely only one of many drivers of genome size in bacteria.

**Fig. 4.**
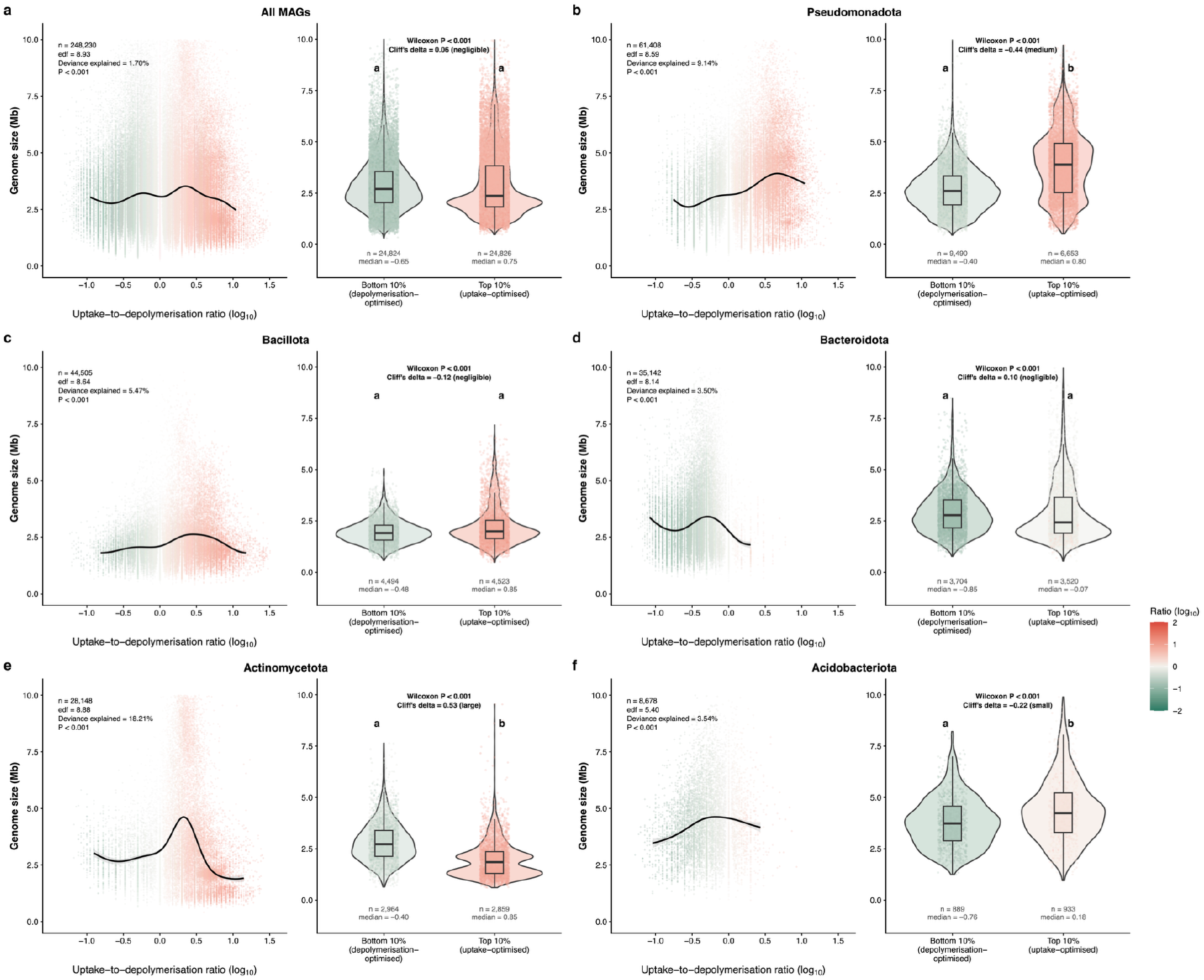
Relationship between genome size and bacterial resource-acquisition strategy. Relationship between genome size and the log_10_-transformed uptake-to-depolymerisation ratio across all bacterial MAGs and the top five bacterial phyla with highest number of MAGs. The left panels show the relationship between genome size and the uptake-to-depolymerisation ratio (log_10_) as evaluated using generalised additive models (GAMs). Points represent individual genomes, coloured by their uptake-to-depolymerisation ratio. The solid line indicates the fitted GAM smooth. Sample size (*n*), estimated degrees of freedom (edf), percentage of deviance explained, and the *P* value of the smooth term are shown. The right panels compare genome sizes of MAGs with the bottom 10% (depolymerisation-optimised) and top 10% (uptake-optimised) of the ratio. Violin plots show the distributions, boxplots show medians and interquartile ranges, and points represent individual genomes. Group sizes and ratio medians are shown below each plot and the colour of violins and boxplots represents the median for each group. Two-sided Wilcoxon rank-sum *P* values and Cliff’s δ are reported. Different letters indicate comparisons that were statistically significant (P < 0.05) and non-negligible in magnitude (|δ| ≥ 0.147).

### Resource acquisition strategies shift during litter decomposition

Next, we set out to experimentally measure variation in the two contrasting resource acquisition strategies identified in bacterial genomic datasets. We used a field experiment to examine the relative prevalence of bacteria assigned to the two strategies during complex plant litter decomposition. Metagenomics was performed on the microbial community growing on litter placed on the soil surface. We hypothesized that uptake-optimised bacteria would be more prevalent at the early stage of decomposition along with fungi who are the pioneers of complex carbohydrate depolymerisation. Indeed, we observed that bacterial MAGs enriched at the early stage of decomposition (2 months since start of litter decomposition experiment) had lower complex carbohydrate depolymerisation genes compared to those enriched at the later stages (7, 14, and 17 months since start of litter decomposition experiment) but showed higher capacity for carbohydrate transport systems (Fig. 5a), indicating the dominance of uptake-optimised bacteria at the early stage. The uptake-to-depolymerisation ratio was highest at the 2-month sampling point and decreased with time (Fig. 5a). The ratio was higher in specific bacterial lineages including Enterobacteriaceae, Pseudomonadaceae, Rhizobiaceae, and Rhodobacteraceae of phylum Pseudomonadota and Micrococcaceae of phylum Actinomycetota all of which were highly enriched at the early stage of decomposition, whereas the bacterial lineages Chitinophagaceae, Sphingobacteriaceae, and Hymenobacteraceae of phylum Bacteroidota and Sphingomonadaceae of phylum Pseudomonadota with lower ratios were highly enriched at the later stage of decomposition (Fig. 5b). These results indicate a temporal shift in bacterial resource acquisition strategies from uptake optimisation during early decomposition towards greater investment in extracellular depolymerisation at later stages, and the lineages specialising in the two strategies are consistent with our global phylogenetic analysis.

**Fig. 5.**
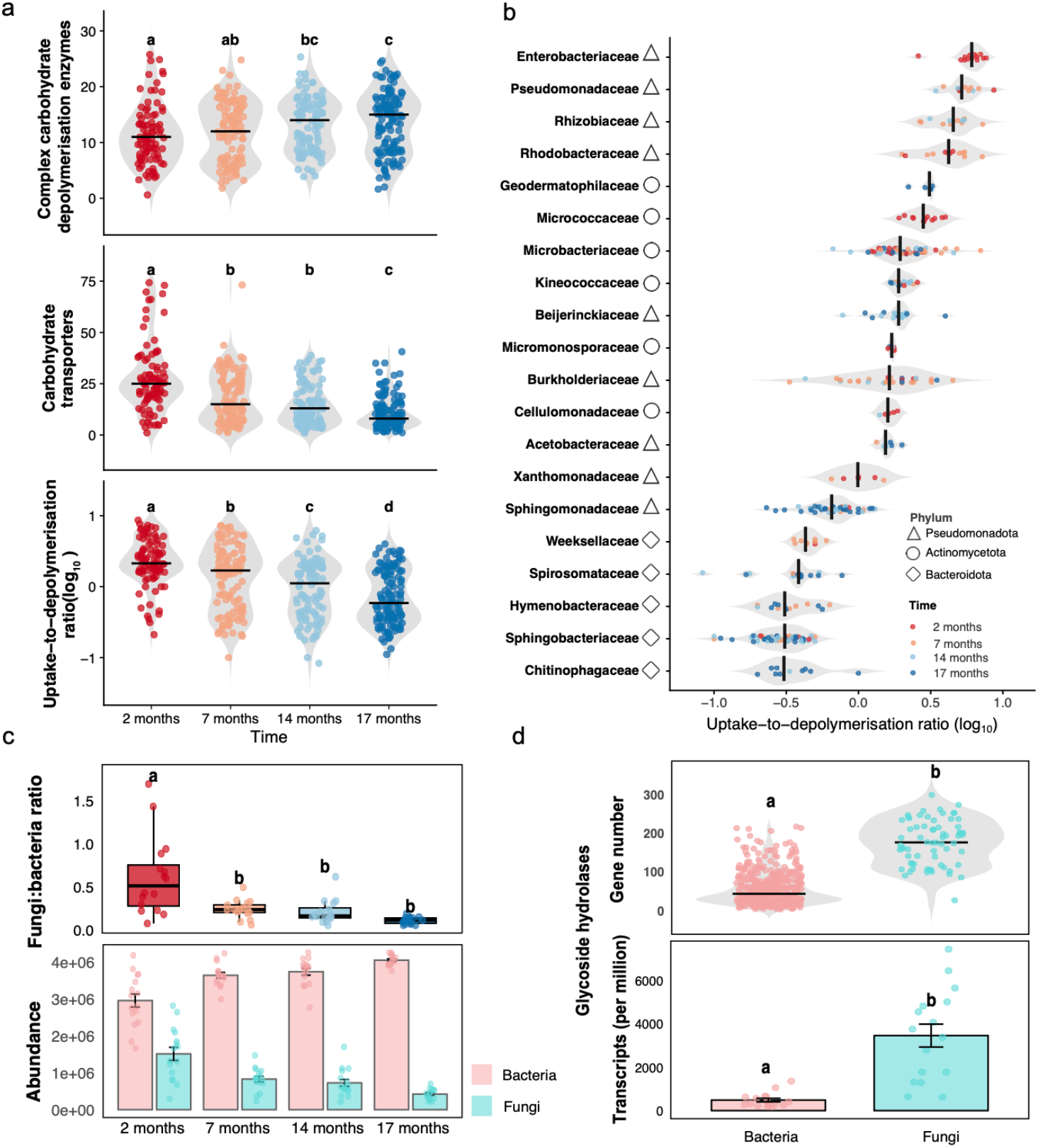
Early litter decomposition is associated with fungal dominance and uptake-optimised bacteria. (a) Bacterial functional repertoires for complex carbohydrate depolymerisation (top), carbohydrate transporters (middle), and the log_10_-transformed uptake-to-depolymerisation ratio (bottom) across four time points: 2, 7, 14 and 17 months after the start of the litter decomposition experiment. Violin plots show distributions with individual MAGs overlaid; black bars indicate medians. Different letters indicate pairwise differences that were both statistically significant (pairwise two-sided Wilcoxon rank-sum tests with Benjamini–Hochberg correction, adjusted *P* < 0.05) and of non-negligible magnitude (|Cliff’s δ| ≥ 0.147, corresponding to at least a small effect). (b) Family-level variation in the uptake-to-depolymerisation ratio across dominant bacterial families with symbols denoting their phyla. Points represent individual MAGs and are coloured by time after the start of the litter decomposition experiment; grey violins show the overall distribution within each family, and black bars indicate medians. (c) Fungi:bacteria abundance ratio (top) and read abundances of bacteria and fungi (bottom) across the four time points. Letters indicate significant differences and effect size of non-negligible magnitude among time points. (d) Glycoside hydrolases (GH) potential and expression by microbial domains. Top, GH gene numbers per MAG of bacteria and fungi. Bottom, domain-level GH transcript abundance in the metatranscriptomes at 2-month time point, partitioned into fungal-and bacterial-encoded GHs. Letters indicate significant differences and effect size of non-negligible magnitude among time points quantified using Cliff’s δ.

Metagenomic analysis of microbial communities at the four time points during litter decomposition suggested greater fungal contribution at the early stages of decomposition, which showed the highest fungal-to-bacterial abundance ratio (Fig. 5c). This temporal pattern is in line with our predictions of fungi as pioneers of plant polymer decomposition. We observed that fungal MAGs contained more extensive capabilities for hydrolytic enzymes than bacterial MAGs and fungal genes for hydrolytic enzymes were strongly expressed compared to bacteria at the early stage of decomposition i.e. 2 months since start of litter decomposition experiment (Fig. 5d). This early-stage dominance of fungi in degrading complex plant polymers in our Mediterranean biome study site is consistent with previous findings from across a wide range of ecosystems^46–48^. Combined with the dominance of uptake-optimised bacteria during early decomposition, these results suggest a functional partitioning in which fungi invest more heavily in extracellular depolymerisation, whereas many bacterial taxa specialise in capturing the resulting soluble products. Such an uptake-optimised strategy may also facilitate microbial cheating, as bacteria — growing alongside fungi on the same litter — can benefit from diffusible compounds released by fungal enzyme activity without making an equivalent investment in extracellular depolymerisation.

An alternative explanation to the dominance of uptake-optimised bacteria during early decomposition is that they preferentially use low-molecular-weight plant compounds that require no depolymerisation. Such plant soluble compounds constitute a transient pool released largely through abiotic leaching^49^. Given the very high fungal investment in extracellular depolymerisation, the early dominance of uptake-optimised bacteria is more parsimoniously explained by the capture of fungal-derived depolymerisation products than by the direct use of low molecular weight plant compounds but explicit separation of mechanisms will likely require isotope based approaches to track the fate of different substrate sources. As decomposition proceeded, bacterial resource acquisition shifted towards greater investment in extracellular depolymerisation. This shift likely reflects changes in litter chemistry, as certain complex plant polymers are degraded and exhausted and the community switches to other substrates^50^. Indeed, CAZyme gene abundances for cellulose declined over the time course, whereas those for hemicellulose and starch increased. In parallel, genes for bacterial peptidoglycan metabolism rose while those for fungal chitin metabolism fell, consistent with an increasing contribution of bacterial relative to fungal necromass at later stages^51^. This temporal pattern suggests that community succession reflects specialisation of different microbial groups on different substrate types.

### A quantifiable proxy of bacterial resource acquisition strategy

Our genome-based approach provides a tractable measure of the relative genomic potential of bacteria for carbohydrate uptake versus complex-substrate depolymerisation, complementing existing trait-based characterisations of microbial resource use^6,52^. A high uptake-to-depolymerisation ratio may arise through several mutually non-exclusive routes: specialisation on low-molecular-weight substrates, residence in labile-carbon-rich niches such as the rhizosphere, or reliance on depolymerisation products released by neighbouring taxa. Although our rhizosphere–bulk soil comparison supports the niche-availability interpretation, resolving the interaction-based route — whether such reliance on others’ depolymerisation products amounts to cheating, which requires a demonstrable fitness cost to the depolymerising taxon — necessitates spatial, co-occurrence and interaction data. Conversely, a low ratio may reflect specialisation on complex polymers, occupancy of niches lacking readily available resources where depolymerisation is the primary route to carbon acquisition, or lifestyles in which breakdown products are privatised rather than released as freely diffusible public goods, for example in biofilms or particle-associated communities^53^, or through specialised cell-surface-associated or transport-coupled enzyme systems^54,55^. Nevertheless, by capturing variation that is phylogenetically conserved, habitat-associated and dynamic during decomposition, the genomic ratio offers a scalable means to compare resource-acquisition strategies across the vast diversity of bacteria.

## Methods

### Collection of publicly available MAGs

We incorporated metagenome-assembled genomes (MAGs) from GlobDB release 226^56^. GlobDB is a species-dereplicated microbial genome resource that integrates genomes from fourteen large-scale catalogues representing environmental and host-associated microbiomes, including marine, freshwater, plant-associated, animal-associated, engineered, and extreme environments. We retained genomes that met minimal quality criteria of at least 50% completeness and less than 10% contamination. After filtering and matching with available functional information, a total of 249,311 MAGs were included in the study. Functional annotations, including CAZyme assignments, KEGG orthologs and Pfam domains, were obtained from the contigs databases and accompanying annotation files provided by GlobDB. This dataset served as a broad comparative reference for bacterial resource acquisition strategies across soil and non-soil systems.

To investigate the environmental distribution of these strategies in soils, we compiled soil MAGs from the Integrated Microbial Genomes and Microbiomes database (IMG/MER)^57^. MAGs annotated under the GOLD ecosystem categories “soil” or “host-associated: rhizosphere” were selected for analysis. We retained only unrestricted genomes that met established quality thresholds, including genomes with at least 50% completeness and less than 10% contamination^58^. After filtering and matching with available functional information, a total of 28,932 soil-associated MAGs were included in the study. All genomes were downloaded on 15 May 2024.

### Trait inference and calculation of the uptake-to-depolymerisation ratio

Functional traits were inferred using the microTrait framework, which maps gene-level functional annotations to trait categories^26^. From the microTrait output, we extracted two sets of traits associated with microbial resource-acquisition strategies. The first captured the potential for complex carbohydrate depolymerisation, defined as the presence of extracellular enzymes involved in the breakdown of major structural polysaccharides, including cellulose, mixed-linkage glucans, xylan and heteroxylan, xyloglucan, heteromannan, and chitin. The second captured carbohydrate uptake capacity, defined as the combined presence of transporter traits responsible for importing low-molecular-weight carbohydrates. For soil MAGs, functional traits were inferred directly using the microTrait framework applied to genome sequences. For GlobDB genomes, trait inference was based on precomputed functional annotations provided by GlobDB, including KEGG orthologs and CAZyme assignments. Specifically, genes corresponding to microTrait-defined depolymerisation and transporter traits were identified by matching KEGG orthologs and CAZy family annotations to the gene sets defined in the microTrait framework. Trait counts derived from GlobDB annotations were therefore directly comparable to those inferred using microTrait, as they relied on the same underlying gene-to-trait mapping rules.

For each genome, transporter-associated and depolymerisation-associated genes were converted to presence–absence profiles and summed within each functional category to estimate transporter and depolymerisation repertoires. We then calculated the uptake-to-depolymerisation ratio by dividing transporter repertoires by depolymerisation repertoires. Log_10_-transformed ratios were used throughout. Each genome was assigned to the high-level habitat category in which it had the highest detection count. The corresponding detection count was used as the analytical weight in both the violin density estimation and statistical analysis. Weighted n for each environmental category was calculated as the sum of detection counts across all genomes assigned to that category and therefore represents the total number of genome detections rather than the number of unique genomes. This meant that habitat-level distributions reflected how frequently each MAG was detected within its dominant habitat, rather than giving rare and frequently detected MAGs equal influence. Differences among environmental categories were assessed using a weighted least-squares model, with the log_10_-transformed uptake-to-depolymerisation ratio as the response variable, habitat category as the predictor and detection count as the weight. Pairwise comparisons of estimated marginal means were performed using heteroscedasticity-robust standard errors with Tukey adjustment for multiple comparisons. Pairwise effect sizes were quantified using a detection-count-weighted form of Cliff’s δ that reflects differences between detection-weighted distributions. Letters from a compact letter display were assigned only where pairwise differences were both statistically significant (Tukey-adjusted *P* < 0.05) and non-negligible in magnitude (|δ| ≥ 0.147), following Romano et al^59^.

### Genome size analyses

Genome size variation was assessed using genomes from the environmental and host-associated MAG dataset. Genome sizes were converted from base pairs to megabases, and genomes with non-finite values or greater than 10 Mb were excluded as per Zhou et al.^60^. Associations between uptake-to-depolymerisation ratio and genome size were evaluated using generalized additive models (GAMs), with genome size as the response variable and the log_10_-transformed uptake-to-depolymerisation ratio fitted as a smooth term. GAMs were first fitted across all genomes and subsequently within major bacterial phyla to assess lineage-specific relationships. For each model, the estimated degrees of freedom (edf), percentage of deviance explained and *P* value of the smooth term were reported. GAMs were fitted using the mgcv package in R^61^.

### Environmental predictors of genomic traits

To examine how environmental factors shape microbial resource-acquisition strategies, we linked genome-level uptake-to-depolymerisation ratios to biomes. Each genome was assigned to a terrestrial biome following the Whittaker classification using mean annual temperature and mean annual precipitation extracted for the sampling location^62^. Geographic coordinates (latitude and longitude) associated with each genome were obtained from the corresponding sampling metadata available in the source databases. For soil MAGs, coordinates were extracted from IMG/GOLD metadata when available. Biome boundaries followed the implementation provided in the plotbiomes R package (https://github.com/valentinitnelav/plotbiomes). Only biomes represented by at least 100 genomes were included in biome-level comparisons. Ratios were log_10_-transformed before statistical analysis. Overall differences in the uptake-to-depolymerisation ratio among biomes were assessed using a Kruskal-Wallis test, followed by Dunn’s pairwise tests with Benjamini-Hochberg correction. Pairwise effect sizes were quantified using Cliff’s δ. Letters from a compact letter display were assigned only where comparisons were both statistically significant (adjusted P < 0.05) and non-negligible in magnitude (|δ| ≥ 0.147)^59^.

To compare resource-acquisition strategies between rhizosphere and bulk-soil communities, we selected IMG/MER MAGs from three sites containing both habitat types: the Populus common gardens at Clatskanie and Corvallis, Oregon, and a Populus cultivar trial in Tennessee. The Clatskanie and Corvallis MAGs were recovered by IMG from the metagenomic datasets reported by Schadt et al. (2024)^30^, whereas the Tennessee MAGs were recovered from the metagenomic dataset analysed by Blair et al. (2018)^31^. MAGs were classified as rhizosphere or bulk soil using the corresponding IMG/GOLD ecosystem metadata. Within each site, differences in the log10-transformed uptake-to-depolymerisation ratio were assessed using two-sided Wilcoxon rank-sum tests, treating MAGs as independent observations, and effect sizes were quantified using Cliff’s δ.

### Phylogenetic structure of resource acquisition strategies

To test whether bacterial resource acquisition strategies exhibited a phylogenetic structure, we used the bacterial reference phylogeny provided in the GlobDB database (release 226). The precomputed species-level tree integrates the standardized GTDB taxonomy and provides a unified backbone phylogeny across all major bacterial lineages. The tree and associated metadata were downloaded directly from the GlobDB resource (https://globdb.org/). Phylogenetic signal in the uptake-to-depolymerisation ratio was quantified as a continuous trait using Pagel’s λ^63^. The phylogeny was pruned to retain taxa with corresponding trait values, and trait values were matched to tree tips. Pagel’s λ was estimated by maximum likelihood, with λ = 0 indicating no phylogenetic signal and λ approaching 1 indicating trait covariance consistent with Brownian evolution. Significance was assessed against a null model with λ fixed at 0. Analyses were performed in R using phytools::phylosig^64^. consenTRAIT was then used to quantify the phylogenetic depth at which the two contrasting resource acquisition strategies were conserved^23^. Because consenTRAIT operates on binary traits, we converted the uptake-to-depolymerisation ratio into two binary traits: uptake-optimised, defined as genomes with top 10% of the ratio, and depolymerisation-optimised, genomes with bottom 10%. The consenTRAIT algorithm identifies the shallowest clades in which at least 90% of descendant taxa share a given trait and computes τD, the mean phylogenetic distance from tips to the most recent common ancestor of these clades. Smaller τD values indicate stronger phylogenetic clustering, whereas larger values indicate phylogenetic dispersion. We implemented this analysis using the getTraitDepth function in the R package castor, with the following parameters: min_fraction = 0.9, count_singletons = TRUE, singleton_resolution = 0, and weighted = FALSE^65,66^.

For visualization, the GlobDB reference phylogeny was imported into R using the ape package^67^ and pruned to retain only genomes present in the GlobDB trait dataset by intersecting tree tip labels with genome identifiers. The pruned tree was visualised using ggtree^68^ with a circular layout. Tip points were coloured by phylum (top 15 most abundant phyla shown explicitly and all remaining phyla grouped as “Other”), and the uptake-to-depolymerisation ratio was overlaid as a single external heatmap ring using ggtreeExtra (geom_fruit)^69^.

### Field site and litter bag experimental design

The litter decomposition experiment was conducted at the Loma Ridge Global Change Experiment site in California, USA, using the same long-term precipitation manipulation plots as described previously^70^. Senescing litter was collected from four replicate plots within each treatment and homogenized. In grassland plots, dry sheath and blade material was cut into ∼10 cm fragments; in shrubland plots, intact fallen leaves were used. Litterbags (15 × 15 cm, 1 mm mesh) were filled with 6 g of air-dried litter and deployed on the soil surface under plant canopies on 12 September 2017. Litterbags were retrieved at four time points 2, 7, 14, and 17 months since start of litter decomposition experiment; 16 bags were sampled at each time point. These time points represented seasonal transitions: 30 November 2017 (end of dry season), 11 April 2018 (end of wet season), 5 November 2018 (end of dry season), and 19 February 2019 (end of wet season).

### DNA extraction and metagenomic sequencing

DNA was extracted from 50 mg of coarse-ground litter samples collected at the four time points using the ZymoBIOMICS DNA Miniprep Kit (Zymo Research, Irvine, CA, USA) according to the manufacturer’s instructions. Samples were homogenized by bead beating for 5 minutes at maximum speed (6.0 m/s) using a FastPrep-24 homogenizer (MP Biomedicals, Irvine, CA, USA). DNA yield and purity were evaluated using agarose gel electrophoresis, a Qubit 3.0 fluorometer (Life Technologies, Carlsbad, CA, USA), and a NanoDrop 2000 spectrophotometer (Thermo Scientific, Waltham, MA, USA). Metagenomic library construction and sequencing were performed at the UC Davis Genome Center (DNA Technologies and Expression Analysis Cores). Libraries were sequenced on the Illumina NovaSeq platform using paired-end 150 bp (PE150) chemistry with an average insert size of 250-400 bp.

### Metagenomic co-assembly, binning, and annotation of prokaryotic and eukaryotic genomes

Metagenomic data processing and analysis were performed using the Metagenome Orchestra (MAGO) pipeline (version V2.2b; 2020-03-08)^71^. For each sample, raw paired-end reads were quality-controlled with FastP (version 0.20.0)^72^, ensuring a minimum read quality of Q30 and removing sequencing adapters. The quality-filtered reads were subsequently assembled using metaSPAdes (version 3.13.0)^73^. Contigs shorter than 1,000 bp were discarded using seqtk (version 1.3-r106) (https://github.com/lh3/seqtk). Taxonomic classification was conducted with Kraken (version 1.1.1)^74^ by querying seqtk-processed metagenomic assemblies against the Kraken database. Resulting taxonomic assignments were aggregated using KrakenTools^75^ to quantify bacterial and fungal abundances. Differences in fungal-to-bacterial abundance ratios across sampling times were tested using one-way ANOVA, followed by Tukey’s Honestly Significant Difference (HSD) test for post hoc multiple comparisons.

Bacterial bins were recovered using the binning module in MetaWRAP (v1.2.1)^76^. High-quality bins (≥50% completeness and ≤10% contamination) were identified using CheckM (v1.0.12)^77^ and taxonomically classified according to the GTDB database^78^. Relative bin abundances of bacterial genomes across samples were quantified using coverM^79^. Temporal differences in complex carbohydrate depolymerisation enzyme repertoires, carbohydrate transporter repertoires and log_10_-transformed uptake-to-depolymerisation ratios were evaluated using pairwise two-sided Wilcoxon rank-sum tests. MAGs recovered at different sampling times were treated as independent observations, and *P* values were adjusted using the Benjamini– Hochberg procedure. Pairwise effect sizes were quantified using Cliff’s δ. Letters from a compact letter display were assigned to sampling times only where comparisons were both statistically significant (adjusted P < 0.05) and non-negligible in magnitude (|δ| ≥ 0.147). For fungal binning, eukaryotic contigs were first identified using EukRep (v0.6.7)^80^. Eukaryotic bins were constructed using MetaBAT2^81^, and gene prediction was carried out with a hybrid pipeline combining GeneMark-ES^82^ and MAKER2^83^. Genome completeness and duplication were assessed with BUSCO (v4.0.6)^84^. Functional annotation of both bacterial and fungal bins was performed by aligning predicted protein sequences to the KEGG database using DIAMOND (v2.0.13)^85^ to assign KEGG orthologs (KOs). Carbohydrate-active enzymes (CAZymes) were identified using dbCAN2^86^. All downstream annotations were parsed using custom scripts to summarize gene functions and metabolic profiles. Analyses focused on glycoside hydrolases (GHs) identified from CAZyme annotations and on transporters for saccharides, polyols, and lipids classified based on KEGG ortholog assignments, as these functional categories capture complementary strategies of extracellular substrate depolymerisation versus transporter-mediated acquisition of low-molecular-weight compounds. Differences in GH gene numbers between bacterial and fungal genomes were assessed using a two-sided Wilcoxon rank-sum test (Mann–Whitney U test), treating genomes as independent observations. Effect size was quantified using Cliff’s δ, with |δ| < 0.147 considered negligible.

### RNA extraction, library preparation, and sequencing

Total RNA was extracted from coarse-ground litter at 2-month sampling point only using the RNeasy PowerSoil Total RNA Kit (Qiagen, Hilden, Germany), following the manufacturer’s instructions. For shrub litter, 0.2 g of material was used, while 0.5 g was used for grass litter. A reduced input mass was used for shrub samples to minimize co-extraction of inhibitory organic compounds and improve RNA yield. After resuspension of RNA in solution SR7, RNA quantity and purity were assessed using a Bioanalyzer 2100 (Agilent, Santa Clara, CA, USA), Qubit fluorometer (Life Technologies, Carlsbad, CA, USA), and NanoDrop 2000 spectrophotometer (Thermo Scientific, Waltham, MA, USA). A total of 200-500 ng of RNA per sample was used for downstream processing. Ribosomal RNA was removed using the Ribo-Zero rRNA Removal Kit (Illumina, San Diego, CA, USA) with a modified protocol combining magnetic beads from yeast and bacterial kits: 0.5× yeast, 0.25× Gram-negative, and 0.25× Gram-positive rRNA removal solutions. Strand-specific, barcoded RNA-seq libraries were prepared using the KAPA RNA HyperPrep Kit (KAPA Biosystems, Cape Town, South Africa) according to the manufacturer’s instructions. Library fragment size distributions were assessed on a Bioanalyzer 2100, and library concentrations were measured using a Qubit fluorometer. Libraries were pooled in equimolar amounts, quantified by qPCR with a KAPA Library Quantification Kit, and sequenced using single-end 100 bp reads on one lane of an Illumina HiSeq 4000 platform. Sequencing was performed at the DNA Technologies and Expression Analysis Cores at the UC Davis Genome Center.

### Metatranscriptomic data processing and functional annotation

Raw metatranscriptomic reads were quality-filtered using Fastp (v0.20.0)^72^ to remove adapter sequences and low-quality bases (Phred score < 30); reads shorter than 50 bp were discarded. Ribosomal RNA (rRNA) sequences were removed using SortMeRNA (v4.3.6)^87^ with comprehensive reference databases, and the remaining non-rRNA reads were retained for downstream analysis. De novo assembly of non-rRNA reads was performed independently for each sample using IDBA-Tran (v1.1.3)^88^. Open reading frames (ORFs) were predicted from assembled contigs using Prodigal (v2.6.3)^89^ in metagenomic mode. All predicted coding sequences were pooled and clustered using CD-HIT-EST (v4.8.1)^90^ at 95% sequence identity to construct a non-redundant unigene catalog. Gene expression quantification was conducted with RSEM (v1.3.1)^91^, using Bowtie2 (v2.4.1)^92^ for alignment. Expression levels were normalized as transcripts per million (TPM) to account for both transcript length and sequencing depth. Taxonomic annotation was performed using Kraken, which classifies reads based on k-mer matching to a reference database, to quantify fungal and bacterial transcripts. Functional annotation was conducted by aligning predicted protein sequences to the KEGG database using DIAMOND (v2.0.13)^85^. Additionally, CAZymes were identified using dbCAN2^86^, enabling classification of glycoside hydrolases, carbohydrate esterases, and other CAZy families. To compare the contribution of fungal and bacterial transcripts to GHs expression, we quantified the GHs expression for each sample and treated observations as independent. Differences between bacterial and fungal groups were assessed using a two-sided Welch’s t-test implemented in the R package rstatix.

## Acknowledgments

We thank Antonio Ribeiro for bioinformatics support; and Claudia Weihe for help with experimental setup and the molecular work. We acknowledge Orange County Parks and the Irvine Ranch Conservancy for field site access. We thank Daan Speth who maintains the GlobDB database. We used ELM, a secure University of Edinburgh-hosted generative AI service (Claude Opus 4.8 model), to assist with language editing and to improve the readability of the manuscript. All AI-assisted text and suggestions were critically reviewed by the authors, who made all scientific decisions and take full responsibility for the final manuscript.

## Author contributions

Conceptualization: AAM, YF

Methodology: AAM, SDA

Investigation: AAM, YF, LCR, KS, JR

Formal analysis: YF

Validation: YF

Data curation: YF

Visualization: YF

Funding acquisition: AAM, JBHM, SDA, YL

Supervision: AAM, YL

Writing – original draft: YF, AAM

Writing – review & editing: All authors

## Data, code, and materials availability

Raw metagenomic sequencing data are publicly available in the NCBI Short Read Archive under BioProject accession PRJNA1178105 with accession numbers from SRR31127223 to SRR31127332: https://www.ncbi.nlm.nih.gov/sra/PRJNA1178105.

Transcriptomics data have been deposited in NCBI’s Gene Expression Omnibus and are accessible through GEO Series accession number GSE148618: https://www.ncbi.nlm.nih.gov/geo/query/acc.cgi?acc=GSE148618.

All processed data are publicly available on figshare under the DOI 10.6084/m9.figshare.31801768. All scripts for data processing and visualization are available via GitHub: https://github.com/MalikLab-UOE/bacterial-uptake-depolymerisation-traits

