## Supplementary figures for "A life-history axis of resource acquisition in bacteria: depolymerisation versus uptake"

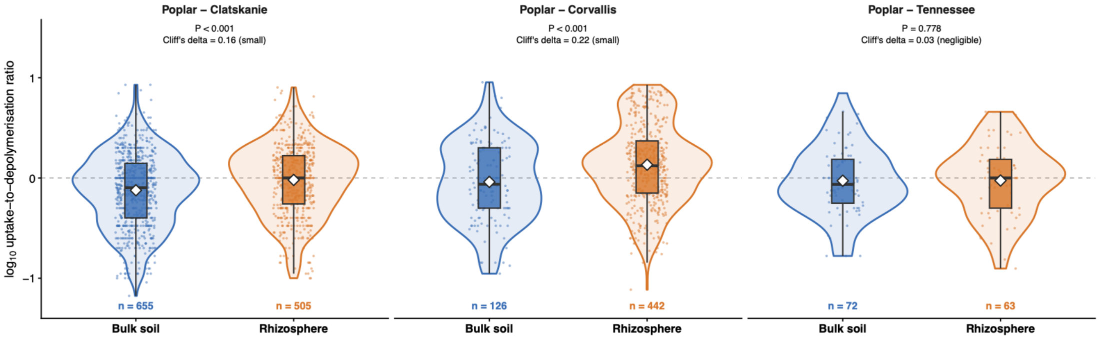


**Figure S1.** Differences in resource-acquisition strategies between bulk and rhizosphere soil bacteria across three field sites of Populus trees. Violin plots show the distributions of $\log_{10}$-transformed uptake-to-depolymerisation ratios for bacterial genomes recovered from bulk and rhizosphere soil samples at Clatskanie, Corvallis and Tennessee sites. Boxplots indicate the median and interquartile range. Pairwise differences between bulk soil and rhizosphere were assessed using two-sided Wilcoxon rank-sum tests, with *P* values shown above each comparison. Effect sizes were quantified using Cliffs $\delta$ and classified as negligible or small. Sample sizes indicate the number of genomes included in each group.

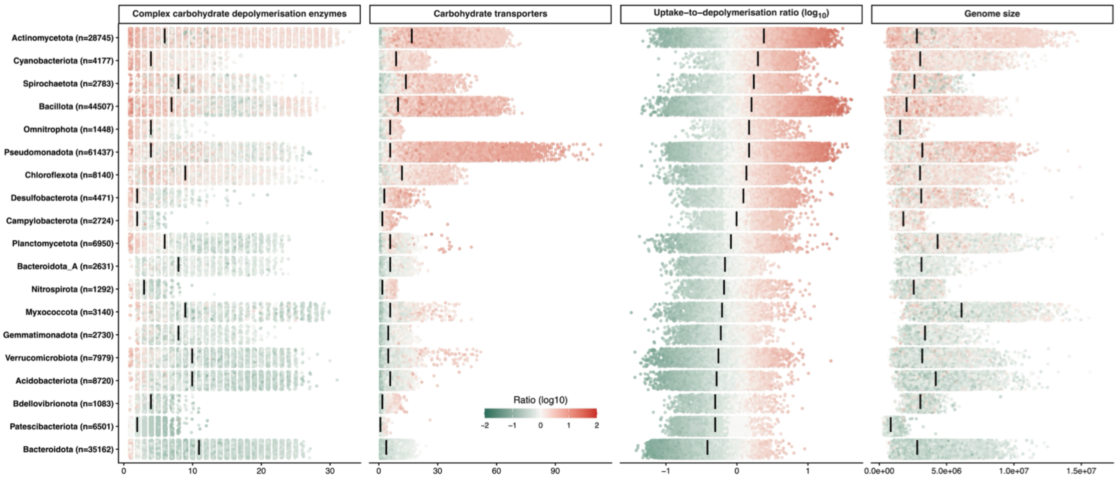


**Figure S2.** Distribution of bacterial resource acquisition strategies across major bacterial phyla. Each point represents a bacterial genome. Distributions of complex carbohydrate depolymerization genes, carbohydrate transporter genes, the $log_{10}$-transformed uptake-to-depolymerization ratio, and genome size are shown for each phylum. Black vertical bars indicate median values.
